# Genetic Architecture and Molecular Networks Underlying Leaf Thickness in Desert-Adapted Tomato *Solanum pennellii*

**DOI:** 10.1101/111005

**Authors:** Viktoriya Coneva, Margaret H. Frank, Maria A. de Luis Balaguer, Mao Li, Rosangela Sozzani, Daniel H. Chitwood

## Abstract

Thicker leaves allow plants to grow in water-limited conditions. However, our understanding of the genetic underpinnings of this highly functional leaf shape trait is poor. We used a custom-built confocal profilometer to directly measure leaf thickness in a set of introgression lines (ILs) derived from the desert tomato species *Solanum pennellii*, and identified quantitative trait loci (QTL). We report evidence of a complex genetic architecture of this trait and roles for both genetic and environmental factors. Several ILs with thick leaves have dramatically elongated palisade mesophyll cells and, in some cases, increased leaf ploidy. We characterized thick ILs 2-5 and 4-3 in detail and found increased mesophyll cell size and leaf ploidy levels, suggesting that endoreduplication underpins leaf thickness in tomato. Next, we queried the transcriptomes and inferred Dynamic Bayesian Networks of gene expression across early leaf ontogeny in these lines to compare the molecular networks that pattern leaf thickness. We show that thick ILs share *S. pennellii*-like expression profiles for putative regulators of cell shape and meristem determinacy, as well as a general signature of cell cycle related gene expression. However, our network data suggest that leaf thickness in these two lines is patterned by at least partially distinct mechanisms. Consistent with this hypothesis, double homozygote lines combining introgression segments from these two ILs show additive phenotypes including thick leaves, higher ploidy levels and larger palisade mesophyll cells. Collectively, these data establish a framework of genetic, anatomical, and molecular mechanisms that pattern leaf thickness in desert-adapted tomato.

## Introduction

Leaves are the primary photosynthetic organs of land plants. Quantitative leaf traits have important connections to their physiological functions, and ultimately, to whole plant productivity and survival. While few aspects of leaf morphology have been unambiguously determined as functional (Nicotra et al., 2011), clear associations between leaf traits and variations in climate have been drawn (Wright et al., 2004). Leaf thickness, the distance between the upper (adaxial) and lower (abaxial) leaf surfaces, has been shown to correlate with environmental variables such as water availability, temperature and light quantity. Thus, on a global scale, across habitats and land plant diversity, plants adapted to arid environments tend to have thicker leaves (Wright et al., 2004; Poorter et al., 2009).

Leaf thickness is a continuous, rather than a categorical, trait. Thus, it is important to distinguish between thickness in the context of “typical” leaf morphology, generally possessing clear dorsiventrality (adaxial/abaxial flattening) in comparison to extremely thick leaves, described as “succulent”, which are often more radial. While the definition of succulence is eco-physiological, rather than morphological (Ogburn and Edwards, 2010), at the cellular level it is broadly associated with increased cell size and relative vacuole volume (Gibson, 1982; von Willert et al., 1992). These cellular traits promote the capacity to store water and to survive in dry environments (Becker, 2007). Allometric studies across land plant families have shown that leaf thickness scales specifically with the size of palisade mesophyll cells - the adaxial layer of photosynthetic cells in leaves (Garnier and Laurent, 1994; Roderick et al., 1999; Sack and Frole, 2006; John et al., 2013). Increased palisade cell height leads to increased area of contact with the intercellular space and thereby to improved uptake of carbon dioxide (CO_2_) into mesophyll cells (Oguchi et al., 2005; Terashima et al., 2011), possibly offsetting the increased CO_2_ diffusion path in thicker leaves. At the organismal level, thicker leaves present a tradeoff between rapid growth versus drought and heat tolerance (Smith et al., 1997). This idea is supported by global correlations between leaf mass per area (LMA), a proxy for leaf thickness, and habits associated with slower growth (Poorter et al., 2009).

Although leaf thickness is a highly functional trait, mechanistic understanding of how it is patterned during leaf ontogeny is poor. The main cellular events that underpin leaf development are the establishment of adaxial/abaxial polarity, followed by cell division, directional expansion, and differentiation (Effroni et al., 2008). Changes in the relative timing (heterochrony) and duration of these events can impact leaf morphology, including thickness. Several mutants have been identified that show clear alterations in leaf thickness. These include the Arabidopsis *angustifolia* and *rotundifolia3* (Tsuge et al., 1996), as well as *argonaute1*, *phantastica*, and *phabulosa* (Bohmert et al., 1998), which have aberrations in the polarity of cell elongation and the establishment of adaxial/abaxial polarity, respectively, as well as the *N. sylvestris fat* and *lam-1* (McHale, 1992, 1993), which affect the extent of periclinal cell division in leaves. However, these developmental mutants do not necessarily inform us of the mechanisms by which natural selection acts to pattern quantitative variation in leaf thickness.

Efforts to understand the genetic basis of leaf thickness in the context of natural variation face several important challenges. First, direct measurement of leaf thickness at a scale that would allow the investigation of Quantitative Trait Loci (QTL) for the trait is not trivial. Because of the difficulty in measuring leaf thickness directly, LMA is most often used as a proxy for this trait (Poorter et al., 2009; Muir et al., 2014). Second, in addition to genetic components, leaf thickness is environmentally plastic – it is responsive to both the quantity and quality of light (Pieruschka and Poorter, 2012). Finally, because leaf thickness varies on a continuous spectrum and is not associated with any particular phylogenetic lineage or growth habit, mechanistic questions regarding its patterning need to be addressed in a taxon-specific manner.

With these considerations in mind, we used two members of the tomato clade (*Solanum* sect. *Lycopersicon*), which are closely related, morphologically distinct, and occupy distinct environments (Nakazato et al., 2010) to study the genetic basis and developmental patterning of leaf thickness. The domesticated tomato species *S.lycopersicum* inhabits a relatively wide geographic range characterized by warm, wet conditions with little seasonal variation. By contrast, the wild species *S. pennellii* is endemic to coastal regions of the Atacama desert of Peru, a habitat characterized by extremely dry conditions (Nakazato, et al., 2010). The leaves of *S. pennellii* plants, therefore, exhibit morphological and anatomical features that are likely adaptations to dry conditions (McDowell et al., 2011; Haliński et al., 2015), including thick leaves (Koenig et al., 2013). Moreover, a set of homozygous introgression lines (ILs) harboring defined, partially overlapping segments of the *S. pennellii* genome in an otherwise *S. lycopersicum* background (Eshed and Zamir 1995) has been used to successfully map a number of QTL, including fruit metabolite concentrations (Fridman et al., 2004; Schauer et al., 2006), yield (Semel et al., 2006), and leaf shape (Chitwood et al., 2013). Here, we used a custom-built dual confocal profilometer to obtain precise measurements of leaf thickness across the IL panel and identified QTL for this trait in tomato. Leaf thickness correlates with other facets of leaf shape, as well as a suite of traits associated with desiccation tolerance and lower productivity. We investigated the anatomical manifestations of thickness in tomato and found a prominent increase in palisade cell height in many thick ILs. Finally, we inferred comparative gene regulatory networks of early leaf development (plastochron stages P1-P4) in two thick lines using organ-specific RNA-Seq and identified molecular networks that pattern *S. pennellii*-like desert-adapted leaves.

## Results

### Complex genetic architecture of leaf thickness across *S. pennellii* ILs

To investigate the genetic architecture and patterning of leaf thickness in the *S. pennellii* IL panel, we used a custom-built dual confocal profilometer device (Fig. S1), which generates precise thickness measurements throughout the leaflet lamina at a range of resolutions (0.1 - 1.0 mm^2^) and at high-throughput. The device makes use of two confocal lasers positioned on either side of the sample and calculates thickness by measuring the distance between each of the sample’s surfaces and the corresponding laser probe. Finally, we visualize thickness as a heatmap of thickness values across the surface of the leaf lamina (Fig. 1A).

**Figure 1.**
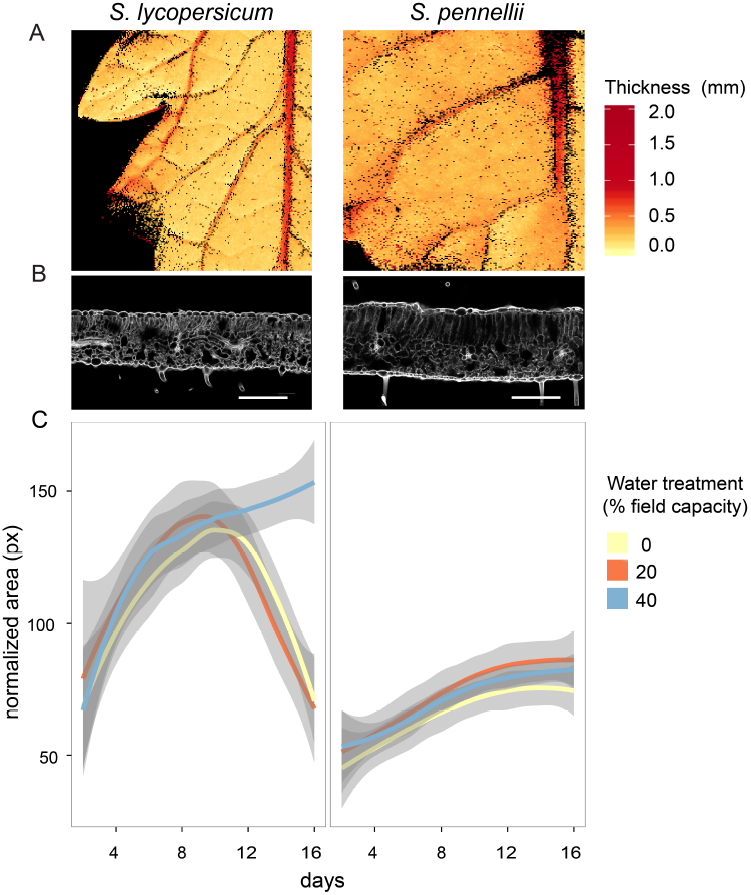
Desert-adapted tomato plants have thicker leaves than domesticated tomato and are resistant to drought. **(A)** Thickness across leaflet blades of domesticated (*S. lycopersicum,* M82) and desert-adapted (*S. pennellii*) tomatoes measured with a custom-built dual confocal profilometer device (Supplemental Figure 1). Median thickness of the *S. lycopersicum* leaflet shown here is 211 μm, and 294 μm for *S. pennellii*. (**B**) Confocal images of propidium iodide stained leaflet cross-sections; scale bar is 200 μm. (**C**) Total shoot area normalized by taking the square root of pixels from top view phenotyping images over 16 days in three water treatments (n=8). Gray shading reflects standard error.

We first compared leaflet thickness in *S. lycopersicum* var. M82 and its desert relative *S. pennellii* LA0716. Our confocal profilometer measurements showed that *S. pennellii* leaflets are thicker than those of domesticated tomato, as previously reported (Fig. 1, Koenig et al., 2013), demonstrating the capacity of this device to quantitatively detect fine differences in leaf lamina thickness. We compared dynamic growth patterns of the two species under water limited conditions and show that, unlike the domesticated species, *S. pennellii* is unaffected by drought (Fig. 1C). This observation highlights the importance of understanding the patterning of developmental traits in this species, such as leaf thickness, which may contribute to drought tolerance. We proceeded to measure leaf thickness across the *S. pennellii* introgression line panel in field conditions.

We used mixed linear regression models to compare each of the introgression lines to the domesticated parent M82 (Dataset S1) and found that 31 ILs had significantly thicker leaflets than the M82 parent, while 5 had transgressively thinner leaflets. The overall broad-sense heritability for leaflet thickness is 39.1% (Fig. 2). The lines with thickest leaflets are IL5-4, IL5-3, IL8-1, IL4-3, IL8-1-1 (contained within IL8-1), and IL2-5, while IL4-1-1, IL2-6-5, IL9-1-3, IL12-4-1, and IL2-1 have thinner leaves than the M82 parent.

**Figure 2.**
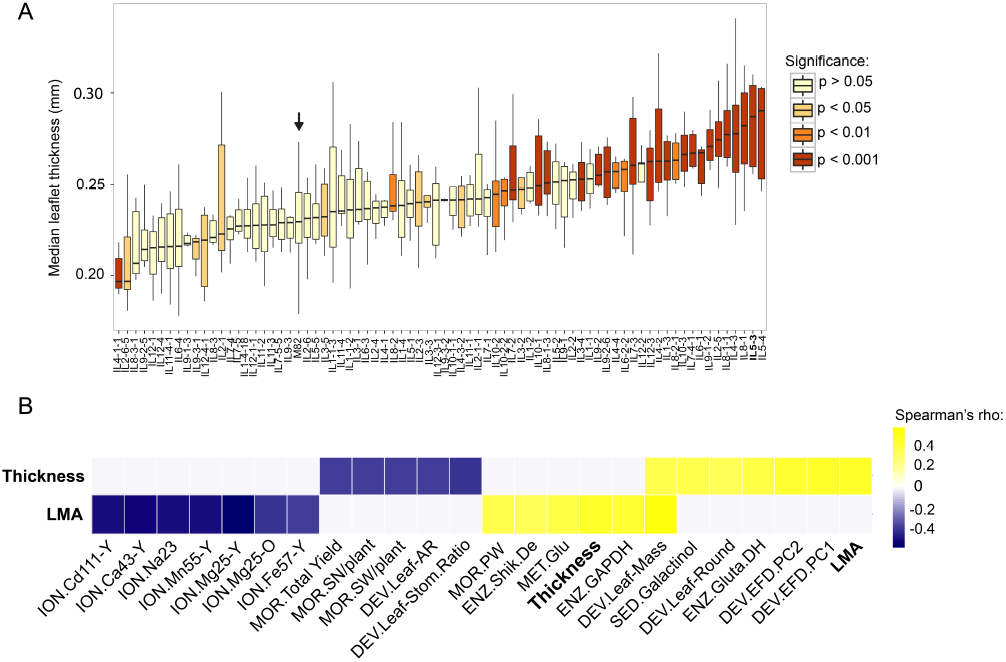
Quantitative Trait Loci for leaf thickness in tomato. **(A)** Leaflet thickness values across the *S. pennellii* introgression line panel. Colors indicate level of significance in comparisons of each IL with M82 (arrow). **(B)** Significant correlations (Spearman’s rho) between leaf thickness (”Thickness”), or leaf mass per area (”LMA”) and a suite of other traits across the *S. pennellii* IL panel (q < 0.05). Traits are grouped by type: ION, elemental profile; MOR, morphological; DEV, developmental; ENZ, enzyme activity; SED, seed metabolite content (Datasets S3 and S4).

Based on the observation that the heritability value for leaf thickness is 39.1 %, we reasoned that environmental factors are likely to play a role in modulating leaf thickness. We thus compared our field experiment with leaf thickness data for vegetative leaves of greenhouse-grown plants. We selected 20 ILs, which were highly significant for leaf thickness differences from M82 in field conditions (p < 0.001) and observed that only some of these lines are also significantly thicker than the domesticated parent in greenhouse conditions (p < 0.05, Fig. S2A). Finally, our observations suggest that leaf thickness varies across the shoot of a number of our select thick leaf ILs with post-flowering leaves having thicker leaves than vegetative leaves (Fig. S2B).

For each leaflet in our field experiment, we also quantified leaf mass per unit area (LMA), which reflects both thickness and density, and is traditionally used as a proxy for leaf thickness. Although the heritability for LMA is similar to that for thickness (33.2% and 39.1%, respectively), significant QTL for these two traits do not consistently overlap (Dataset S1).

### Leaf thickness and LMA are correlated with distinct suites of traits in tomato

We generated pairwise correlations between leaflet thickness, LMA, and a suite of other previously published traits including metabolite (MET), morphological (MOR), enzymatic activity in fruit pericarp (ENZ), seed-related (SED), developmental (DEV), and elemental profile-related (ION) (Datasets S2-4, Chitwood et al., 2013 and references therein). Spearman’s correlation coefficients with significant q-values (q < 0.050) are reported in Fig.2B. Leaf thickness and LMA are correlated (rho = 0.423, q = 0.003). Leaf thickness also correlates with leaf shape parameters, such as roundness (rho = 0.328, q =0.044), aspect ratio (rho = -0.327, q = 0.045), and the first two principal components of the elliptical Fourier descriptors of leaflet shape (EFD.PC1 rho = 0.414, q = 0.004 and EFD.PC2 rho = 0.406, q = 0.005). Thickness is negatively correlated with several reproductive traits, including yield (rho = -0.337, q = 0.037), seed weight (rho = -0.342, q = 0.033) and seed number per plant (rho = -0.339, q = 0.036). Moreover, leaf thickness is negatively correlated with leaf stomatal ratio, the relative density of stomata on the abaxial and adaxial sides of the leaf (rho=-0.352, q = 0.031), and positively with glutamate dehydrogenase activity (rho = 0.367, q = 0.017) and seed galactinol content (rho = 0.342, q = 0.048).

Leaf mass per area is associated with a distinct suite of traits from leaf thickness. In addition to a positive correlation with the content of some enzymes (GAPDH and Shikimate DE) and metabolites (Glutamate), LMA is significantly negatively correlated with the accumulation of Na and Mg in all leaflets tested. LMA, but not leaf thickness, is also significantly positively correlated with total plant weight, reflecting vegetative biomass accumulation.

### Thick IL leaves have elongated palisade parenchyma cells

Leaf cross-sections of field-grown M82 and select ILs with increased leaf thickness, as well as greenhouse-grown *S. pennellii* (Sp) leaves were stained with propidium iodide to assess the anatomical changes that lead to increased leaf thickness. We observed that, relative to the M82 parent, the Sp parent and several ILs, have an elongated palisade mesophyll cell layer corresponding to the adaxial layer of photosynthesizing cells in tomato leaves (Fig. 3). Palisade parenchyma elongation is especially dramatic for IL1-3, IL2-5, IL4-3, and IL10-3. Both leaf thickness and palisade elongation phenotypes are attenuated for vegetative leaves of greenhouse-grown plants (Fig. S2, Fig. S3A).

**Figure 3.**
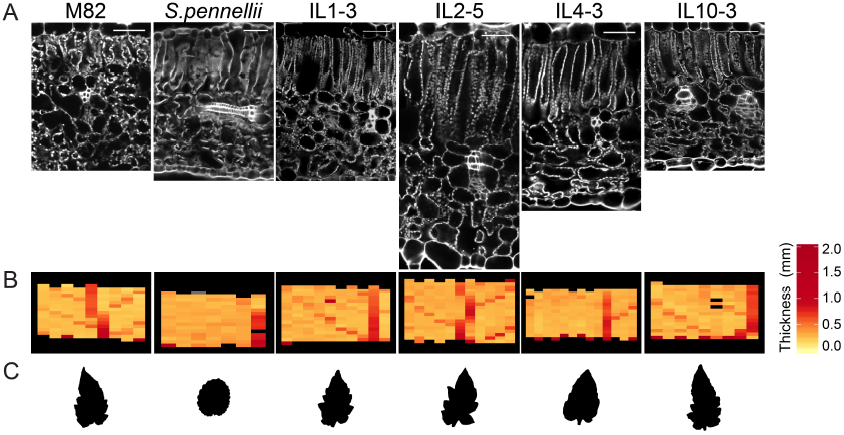
Anatomical manifestations of thicker leaves. **(A)** Confocal images of propidium iodide stained cross-sections of field-grown M82, select ILs and S. pennellii grown in greenhouse conditions; scale bars are 50 μm**. (B)** Representative leaf thickness plots and (C) leaflet binary images of field-grown plants as for (A).

### Anatomy and early leaf development in select ILs with thick leaves

To capture an overall view into the core mechanisms of leaf thickness patterning, we further analyzed lines IL2-5 and IL4-3. We selected IL2-5 due to its dramatic anatomy in field conditions (Fig. 3) and its lack of other characterized leaf morphology phenotypes (Chitwood et al., 2013), while IL4-3 leaflets are both significantly thicker and less serrated than those of the domesticated parent (Fig. 2; Dataset S1, circularity - the ratio between leaflet area and the square of its perimeter - reflects lobing and serration). To further investigate the relationships between genetic determinants of leaf thickness in these ILs, we generated a double homozygous line combining the entire *S. pennellii* segments of IL2-5 and IL4-3.

Double homozygotes (IL2-5/IL4-3) have significantly thicker leaves than M82 at both vegetative (Fig. 4A, p = 0.019) and post-flowering stages (Fig. S2B) in greenhouse conditions. Additionally, IL2-5/IL4-3 plants have significantly smoother margins than either of the IL parents (Fig. 4B), suggesting additive genetic interactions for both of these traits. We next compared the dimensions of the mesophyll cell layers in each IL and the double homozygote line to determine the contributions each cell layer makes to the observed increase in leaflet thickness. We found that palisade mesophyll cells are significantly larger in IL2-5/IL4-3 than in M82 leaves (Fig. S4). Further, the ratios of palisade cell length to both total leaf thickness and to the length of the spongy mesophyll are significantly larger in IL2-5/IL4-3 than in M82 leaves (Fig. S4). IL2-5 shows similar albeit less pronounced trends as the double homozygote line, while in IL4-3 both spongy and palisade mesophyll cell layers are longer than in M82, with the spongy mesophyll layer making the most significant contribution to leaf thickness.

**Figure 4.**
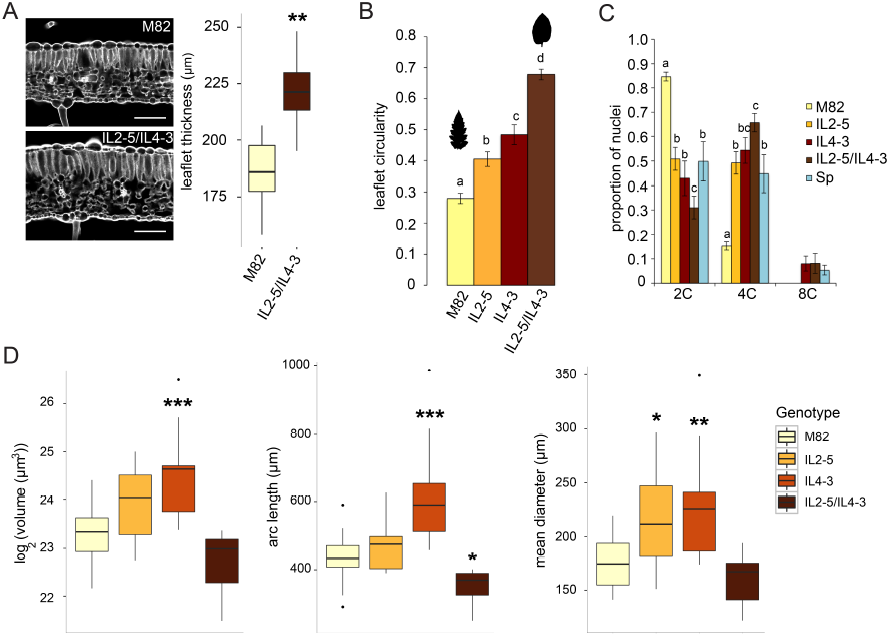
Leaf morphology and ploidy of IL2-5/IL4-3 double homozygote plants. **(A)** Representative propidium iodide-stained leaflet cross-sections (left) and thickness measurements (right) for the 7th leaf of greenhouse-grown M82 and double homozygous IL2-5/IL4-3 plants (n = 10). Scale bars are 200 μm; ^**^ p < 0.01. **(B)** Circularity (ratio of area to the square of the perimeter) of distal lateral leaflets as in (A). Silhouettes of representative M82 and IL2-5/IL4-3 leaflets are shown above bars. Letters indicate statistical significance in each pairwise genotype comparison (p < 0.05). **(C)** Distribution of relative nuclear sizes reflecting endoreduplication in leaflets as in (A) and (B) (n = 5). Letters denote statistical significance between pairwise genotype comparisons at each ploidy level. (D) Leaf plastochron P3 dimensions calculated from 3D surface reconstructions of vegetative shoot apices (n = 9; ^*^ p < 0.05, ^**^ p < 0.01 relative to M82).

Since increases in cell size are often driven by endopolyploidy, we performed flow cytometry on fully expanded vegetative leaves of each genotype and observed increased ploidy profiles in all lines relative to the domesticated parent (Fig. 4C). Notably, the double homozygote line exhibited higher ploidy levels than both single ILs and the *S. penellii* parent (Fig. 4C, Fig. S4). Notably, we also observed a trend to increased ploidy in several greenhouse-grown thick ILs (IL7-4-1, IL8-1) (Fig. S3B).

To understand if alterations in leaf size occur during early stages of leaf ontogeny in these lines, we quantified P3 organ dimensions and compared them with the M82 parental line. For this, we assembled 3D confocal reconstructions of vegetative shoot apices, calculated the surface mesh, extracted P3 leaf primordia, and quantified their total volume, length, and mean diameter. We found that IL4-3 P3 leaf primordia are significantly larger than M82 in terms of overall volume (p = 0.0179), as well as both length (p = 0.0035) and diameter (p = 0.0230). In IL2-5 P3 volume (not statistically significant) and diameter (p = 0.0116) are increased, while length is comparable to M82. Although P3 primordia of double homozygote plants were statistically indistinguishable from those of M82 plants except for shorter arc length (p = 0.0411) (Fig. 4D) our observations also suggest that double homozygote leaves increase in size dramatically between P3 and P4 stages (Fig. S5).

### Transcriptomic signatures of early leaf development in thicker ILs

To investigate the molecular events that define the patterning of IL2-5 and IL4-3 leaves, we isolated leaf primordia from each IL and the two parents (M82 or Sl and Sp) at four successive stages of development: P1 (containing the shoot apical meristem, SAM, and the youngest leaf primordium), P2, P3 (characterized by leaflet emergence) and P4 (typically the onset of cell differentiation) (Fig. 5A). For *S. pennellii*, P1 samples were comprised of the SAM, P1, and P2, since these organs were not separable by hand dissection. Thus, the Sp transcriptomic dataset includes samples designated as P1, P3, and P4. Principal Component Analysis (PCA) of the resulting RNA-Seq data, after normalization and filtering, shows that samples group clearly by organ stage (PC2) (Fig. 5B). In addition, PC1 separates *S. pennellii* samples from all other genotypes. To investigate how IL leaves are similar to the Sp parent, we looked for genes that are differentially expressed (DEGs) between corresponding stages of each IL and the M82 parent, while also being differentially expressed between M82 and Sp. In other words, we identified the set of DEG for each organ stage that is common to each IL and Sp relative to M82. For P2 we considered only the comparison with M82, as our Sp dataset did not include independently dissected P2 stage primordium samples (Fig. S6, Dataset S5).

**Figure 5.**
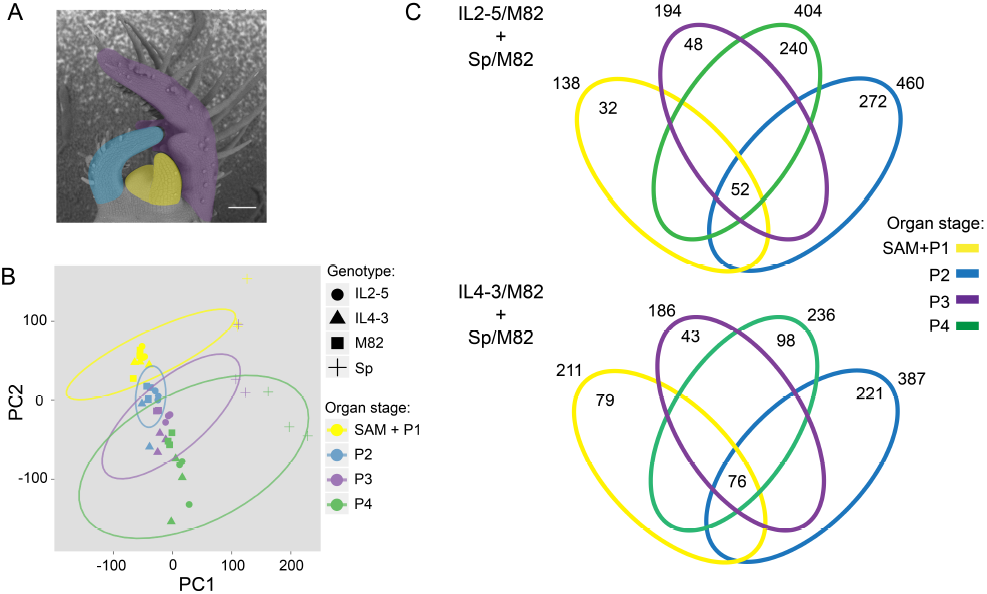
Comparative transcriptomics of leaf development in two thick ILs and their parents. **(A)** Successive stages of leaf development (plastochrons P1-P4 colored as in legend in (B)) were dissected from M82, *S. pennellii* (Sp) and thick ILs 2-5 and 4-3. **(B)** Principal Components Analysis (PCA) of normalized RNA-Seq read counts. **(C)** Venn diagrams (not to scale) depict an overview of differentially expressed genes (DEGs, q < 0.05) that are shared in each IL and the Sp parent relative to M82. The number of DEGs unique to each organ is shown within elipses and those common to all organs, in the center. The total number of DEGs at each plastochron stage is shown outside ellipses.

We identified a total of 812 DEGs across P1-P4 stages in IL2-5, and of these, 544 are up-regulated in at least one organ stage, while 269 are down-regulated (Fig.5C). In IL4-3, we detected 632 DEG, 361 of which are up-regulated and 271 are down-regulated in the IL (Fig. 5C). Many of the DEGs are differentially expressed at more than one stage (Fig. 5C, Dataset S5). Additionally, based on tomato transcription factor (TF) annotation by Suresh et al. (2014), we identified putative transcription factor-encoding genes among each IL’s DEG sets. Myb-related, Ethylene Responsive, MADS, and WRKY are the abundant classes of TF-encoding DEGs in IL2-5, while in IL4-3 TFs belonging to bZIP and Myb-related are highly represented families (Fig. S7).

We identified differentially expressed TF-encoding genes that are common to the two ILs and the Sp parent (Fig. 6), reasoning that some of these can be regulators of leaf thickness. Five of the seven shared TF-encoding genes are up-regulated in the ILs relative to M82. A MADS-box TF (Solyc12g087830) is up-regulated at all stages in both ILs, while two additional inflorescence meristem-related transcription factors, LFY-like (Solyc03g118160) and AP2-like (Solyc07g049490) are differentially expressed at corresponding stages in both ILs. The SHORTROOT-like (SHR-like) GRAS TF Solyc08g014030 is up-regulated at P2 in both ILs, while its expression increases at each progressive stage and peaks at P4 in all genotypes. A putative JASMONATE ZIM-domain protein (JAZ1, Solyc12g009220) is also up-regulated at P2 in both ILs, while a LIM-domain protein (Solyc04g077780) is up-regulated in the ILs at P3 (in IL4-3) and P4 (both ILs) (Fig. 6A).

**Figure 6.**
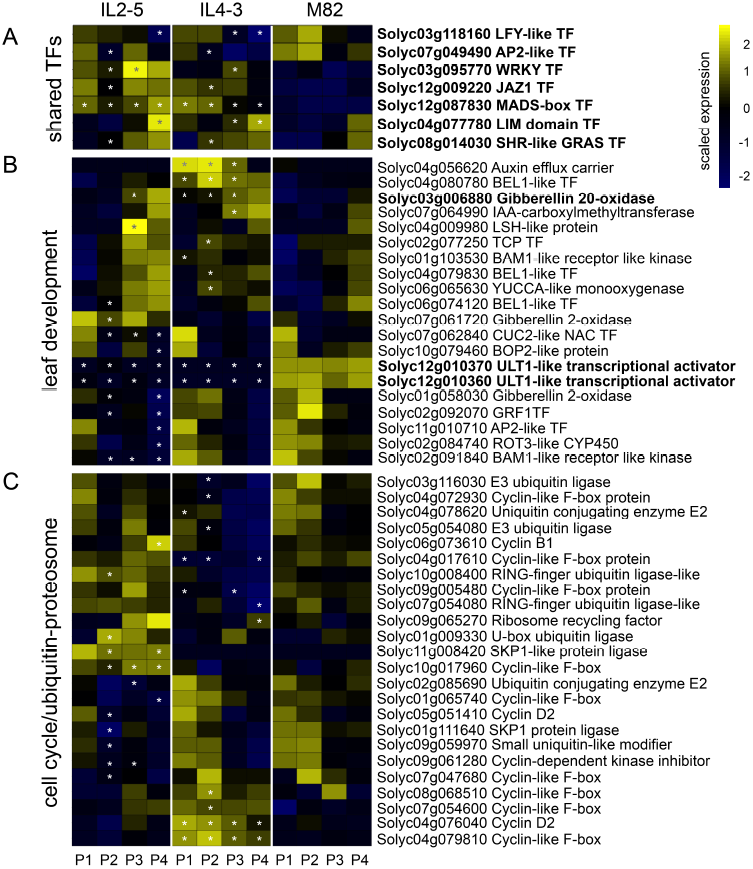
Comparative expression profiles of genes in three functional categories across leaf development (P1 – P4) in thick ILs 2-5 and 4-3: **(A)** Transcription factors common to both ILs. **(B)** Genes involved in leaf development in tomato (as in Ichihashi et al., 2014), and **(C)** Gene annotated to encode components of the cell cycle or ubiquitin protesaome pathway (contain one of the terms “cell cycle”, “cyclin”, “ubiquitin”, “E2F”, “mitosis”, “mitotic”, “SKP”). Plastochron stages with statistically significant DE (q < 0.05) relative to M82 are marked with an asterisk. Genes, which are differentially expressed in at least one stage in both ILs are marked in bold.

Next, we compared the expression profiles of genes known to be involved in tomato leaf development (Ichihashi et al., 2014). We selected only genes that are differentially expressed in the same direction in each IL and Sp relative to the domesticated parent M82 and highlighted genes that are common to both thick ILs to arrive at a set of entities that may be core to the patterning of leaf thickness (Fig. 6B). A gibberellin 20-oxidase encoding gene (GA 20-ox, Solyc03g006880) is up-regulated at P3 in both ILs and throughout the P1-P3 interval in IL4-3. A set of two closely related ULTRAPETALA1 genes (ULT1, Solyc12g010360 and Solyc12g010370) is down-regulated at all leaf developmental stages in both ILs. A number of leaf development regulators are additionally differentially expressed in either of the ILs. Some noteworthy classes include entities related to auxin metabolism or transport (auxin efflux carrier, IAA-carboxymethyltransferase, YUCCA-like monooxygenase), leaf complexity, lobing and serrations (three BEL1-like TFs, CUC2-like and BOP2-like), meristem maintenance or patterning (two BAM1-like receptor kinases and an AP2-like TF), and cell division and expansion (GRF1 and ROT3-like TFs).

Similarly, we also queried DEG sets for entities annotated as cell cycle or endoreduplication to assess whether these two thick ILs share a common trajectory of cellular events during leaf ontogeny (Fig. 6C). Overall, we observed distinct expression profiles for these genes in IL2-5 and IL4-3.

Finally, to broadly characterize the types of processes that may regulate the molecular networks of early leaf development in the ILs we applied GO enrichment analysis (agriGO, Du et al., 2010) (Dataset S6) and identified statistically enriched promoter motifs among the organ-specific DEG sets (Dataset S7). Importantly, we observed that at P4, the set of up-regulated genes in IL2-5 is enriched for biological process terms relating to “photosynthesis” (GO:0015979) and “translation” (GO:0006412), while down-regulated genes at this stage are enriched for terms relating to “DNA binding” (GO:0003677). Our promoter motif analysis showed that motifs associated with regulation by abiotic factors such as light, circadian clock, water availability, and temperature are prominent among IL2-5 DE genes. In addition, binding sites for developmental regulators, hormone-associated promoter motifs, and a cell cycle regulator are among the list of significant motifs. Among development-associated motifs, CArG (MADS-box), BEL1-like (BELL) and SBP-box transcription factor binding sites are also significantly enriched in both IL 2-5 and 4-3 DEG sets. (Fig. S8, Dataset S7).

### Gene regulatory networks of early leaf development in thick ILs

To detect regulators of early leaf development that each IL (IL2-5 and IL4-3) shares with the *S. pennellii* parent, we inferred Dynamic Bayesian Networks (DBN) using the IL and Sp overlapping DEG sets described in the previous section (de Luis Balaguer et al., 2017). Additionally, we only allowed putative transcription factor-encoding genes (Suresh et al., 2014) as “source” nodes (genes that control the expression of other co-expressed genes). First, we constructed individual networks for each leaf developmental stage, for which an overlap with Sp data is available (P1, P3, P4), and then combined the results to visualize the overall *S. pennellii*-like leaf developmental networks (Fig. 7, Dataset S8). The IL2-5 network (Fig. 7A) contains two major regulators, which are central to more than one developmental stage: a SQUAMOSA promoter-binding protein-like domain gene (SBP-box 04g, Solyc04g064470) and a CONSTANS-like Zinc finger (Zn-finger CO-like 05g, Solyc05g009310) (Dataset S8). Similarly, the IL4-3 network (Fig. 7C) features two central regulators: a BEL1-like homeodomain transcription factor gene (BEL1 04g, Solyc04g080780) and a MADS-box domain-containing gene (MADS-box 12g, Solyc12g087830) (Dataset S8). Importantly, few nodes are shared between the organ-specific networks of IL2-5 and IL4-3. We surveyed each network for shared differentially expressed leaf development genes and found that GA 20-ox 03g (Solyc03g006880) is present in both networks but is regulated by different sets of transcription factors in each IL (Fig. 7B, D).

**Figure 7.**
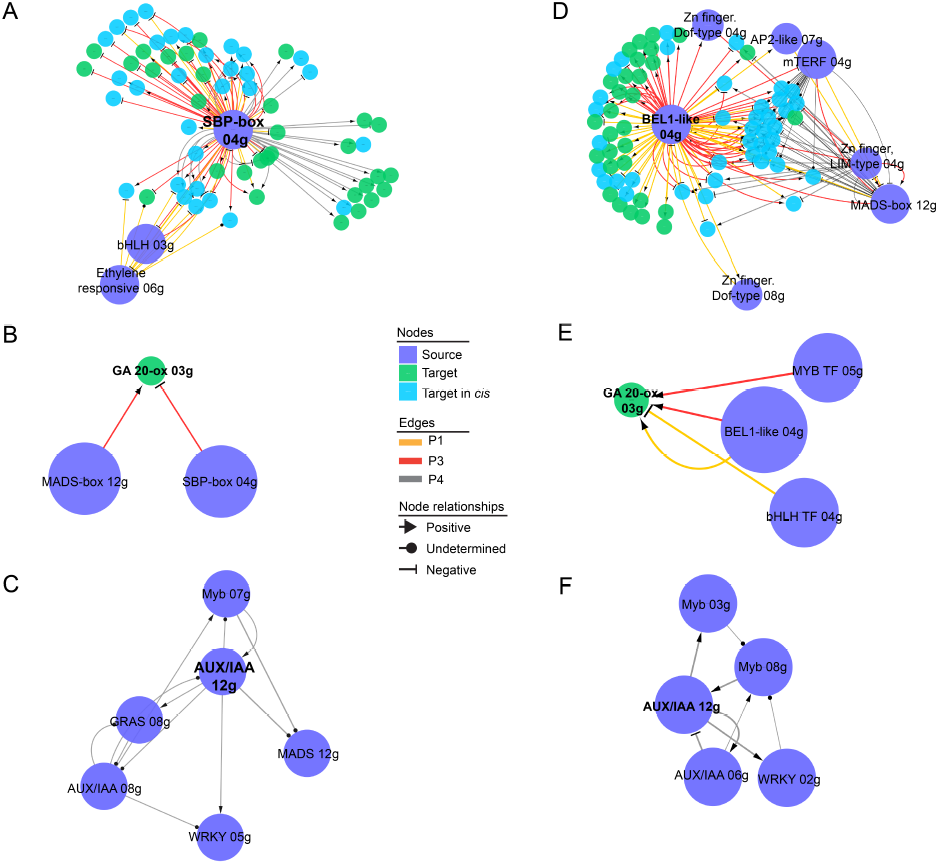
Select leaf development gene regulatory sub-networks for (A-C) IL2-5 and (D-F) IL4-3. Sub-networks for regulators central to more than one plastochron stage are shown in **(A)** and **(D)**. A GA 20-oxidase gene (GA 20-ox 03g, Solyc03g006880) and its regulators in each IL **(B)** and **(E)**. Sub-networks of dynamic gene regulatory networks, showing interactions of an AUX/IAA TF (AUX/IAA 12g, Solyc12g096980) with other source nodes **(C)** and **(F)**. Gene IDs of highlighted nodes: SBP-box 04g, Solyc04g064470; BEL1-like 04g, Solyc04g080780; MADS-box 12g, Solyc12g087820; MYB TF 05g, Solyc05g007710; bHLH TF 04g, Solyc04g074810; GRAS 08g, Solyc08g014030; Myb 07g, Solyc07g052490; WRKY 05g, Solyc05g015850; AUX/IAA 06g, Solyc06g008580; Myb 03g, Solyc03g005570; Myb 08g, Soly08g005870; WRKY 02g, Solyc02g080890. Nodes and edges are colored according to legend.

We also inferred a second set of networks for each of the ILs by identifying DEGs using similar criteria as above. However, in contrast to the previous set of networks, where genes were separated into organ stages based on differential expression at each discrete stage, we used a clustering approach to group regulators and select co-expressed gene sets according to expression profiles. For these analyses, we also included P2 DEGs (IL vs M82) to ensure continuity of expression profiles (Dataset S9). This approach allowed us to examine a more dynamic view of early developmental processes. The resulting networks (Dataset S9) feature a putative auxin responsive TF AUX/IAA 12g (Solyc12g096980) for both ILs (Fig. 7E, F). Moreover, the AUX/IAA 12g sub-network or IL2-5 includes the SHR-like GRAS domain TF that is up-regulated during leaf development in both ILs (GRAS 08g, Solyc08g014030) (Fig. 6A, Fig. 7E).

## Discussion

### Leaf thickness has a complex genetic architecture in desert-adapted tomato and is associated with overall leaf shape, desiccation tolerance, and decreased yield

While extensive progress has been made dissecting the molecular-genetic patterning of two-dimensional leaf morphology, relatively little is known about the third dimension of leaf shape – thickness. Here, we used a custom-built dual confocal profilometer to obtain direct measurements of leaf thickness across the *S. pennellii* x *S. lycopersicum* IL panel (Eshed and Zamir, 1995) (Fig. 1, Fig. S1) and identified QTL for this trait (Fig. 2A). We found that nearly half of the ILs have significantly thicker leaves than the domesticated parent M82, while a small number have transgressively thinner leaves. The broad-sense heritability for leaf thickness in this experiment is moderate (39%). Collectively, these observations point to a complex genetic basis for this trait. A previous quantitative genetic analysis of a suite of desert-adaptive traits in the same *S. pennellii* IL panel found fewer significantly thicker lines and lower heritability (12%) for this trait (Muir et al., 2014). However, the previous study estimated thickness as the ratio of LMA to leaflet dry matter content, while we measured thickness directly. Further, our study was conducted in field conditions, while Muir et al. (2014) measured the trait using greenhouse-grown plants. Given that environment significantly affects the magnitude of this trait (Fig. S2) it is not surprising that these studies report only partially overlapping outcomes.

In order to understand how variation in leaf thickness relates to other traits, particularly to leaf mass per area, we calculated pairwise correlation coefficients among all leaf shape and elemental profile traits, as well as a collection of previously published traits (summarized in Chitwood et al., 2013; Datasets S3, S4). As expected, leaf thickness and LMA are significantly correlated across the IL panel. However, the two traits have distinct sets of significant trait correlations (Fig. 2B). Collectively, these data suggest that thickness and LMA are likely patterned by separate mechanisms and that direct measurements of leaf thickness are necessary to further dissect the genetic basis of this trait.

Leaf thickness is significantly correlated with leaf shape traits such as aspect ratio and the first two principal components of elliptical Fourier descriptors of overall shape. However, our data do not establish whether this correlation reflects a common patterning mechanism or developmental and/or mechanical constraints among these traits. Alternatively, the relatively modest correlations (rho values between 0.33 - 0.41) could reflect independent variation in these traits resulting in considerable flexibility in final leaf morphology, as suggested by Muir et al. (2016).

Leaf thickness is negatively correlated with yield-related traits, which suggests a trade-off between investments in vegetative and reproductive biomass that is further substantiated by the positive correlation between LMA and plant weight (Fig.2B). Some studies support the hypothesis of a tradeoff between leaf mass per area and rapid growth (Smith et al., 1997; Poorter et al., 2009), while others find poor coordination between growth rate and LMA (Muir et al., 2016). Finally, leaf thickness is significantly correlated with leaf stomatal ratio, glutamate dehydrogenase activity, and galactinol content in seeds, a suite of traits associated with desiccation tolerance in plants (Taji et al., 2002; Lightfoot et al., 2007). We also observed negative correlations between LMA and the accumulation of several elements in leaves, most notably Na and Mg (Fig. 2C). This finding supports the idea that LMA and thickness are distinct traits, and that LMA reflects the material composition of leaves, while leaf thickness is a developmentally patterned trait.

### Thicker *S. pennellii* IL leaves have elongated palisade mesophyll cells

The observed elongated palisade mesophyll cells in the leaves of several field-grown ILs with significantly thicker leaves (Fig. 3A), as well as in the desert-adapted *S. pennellii* parent suggest that dorsiventral expansion of palisade mesophyll cells contributes most prominently to increased leaf thickness. This hypothesis is supported by the fact that palisade cell height increases more significantly than the total height of the spongy mesophyll in thick leaves of double homozygous IL2-5/IL4-3 lines (Fig. S4). Palisade cell height is positively correlated with photosynthetic efficiency (Niinemets et al., 2009; Terashima et al., 2011) and water storage capacity in succulent CAM (Crassulacean Acid Metabolism) plants (Nelson et al., 2005). Our data also indicate that the magnitudes of palisade cell elongation, as well as overall leaf thickness are modulated by environmental inputs (Fig. 2, Fig. S2). High light has been shown to mediate increased leaf thickness (Poorter et al., 2009; Wuyts et al., 2012; Kalve et al., 2014), as well as specifically palisade cell elongation (Kozuka et al., 2011) in Arabidopsis, while elongated palisade cells promote a more efficient distribution of direct light throughout the photosynthetic mesophyll compared with shorter cells (Brodersen et al., 2008; Brodersen and Vogelman 2010). Thus, thicker leaves composed of elongated palisade cells may be an adaptation to desert-like dry, direct light environments, whereby the magnitude of these traits is responsive to these environmental cues. Consistent with this hypothesis, we observed that IL2-5 DEG promoters are enriched in motifs that reflect sensitivity to abiotic stimuli, prominently light and water status (Fig. S8, Dataset S7).

### Mechanisms of cell enlargement in thick ILs: increased ploidy and alterations in cell cycle related gene expression

We compared the size of palisade mesophyll cells in leaf cross sections of thick ILs 2-5, 4-3, and a homozygous line combining both introgression segments and observed larger palisade cells compared to M82 (Fig. S4), suggesting a link between leaf thickness and cell size in tomato. Further, we showed significantly higher ploidy levels in the leaves of these lines relative to the domesticated parent (Fig. 4C), indicating that increased endoreduplication may underpin larger cells, and ultimately, thicker leaves. A partially overlapping series of cell division, cell expansion, and cell differentiation events underlie leaf development (Effroni et al., 2008). These processes are tightly coordinated to buffer perturbations in overall organ shape and size (Tsukaya 2003; Beemster et al., 2003). Thus, the relative timing and duration of any of these events can impact leaf size and morphology. Additionally, different tissue types in the leaf can have distinct schedules of cellular events during leaf ontogeny; for example, in Arabidopsis palisade mesophyll cells have a shorter window of cell division compared to epidermal cells, and thus an earlier onset of cell expansion and endoreduplication, resulting in differences in cell volumes and geometry (Wuyts et al., 2012; Kalve et al., 2014). Given the prominent contribution of specific cell types to leaf thickness (palisade mesophyll cells in IL2-5, for example, vs both palisade and spongy mesophyll cells in IL4-3 (Fig. S4)), kinematic studies to capture the timing and extent of tissue-specific cell division and endoreduplication are needed to fully address the dynamic cellular basis of leaf thickness patterning. The observed increase in P3 organ volume and thickness in IL4-3 and, to a lesser extent, IL2-5 relative to M82 (Fig. 4D) support the notion that differences in the trajectory of cellular events during early leaf ontogeny may underpin leaf thickness.

Comparative gene expression profiles of early leaf ontogeny in ILs 2-5 and 4-3 show evidence of *S. pennellii*-like alterations in cell proliferative activity in these thick ILs. Specifically, among a small set of shared differentially expressed genes is a GRAS-domain TF GRAS 08g (Solyc08g014030) up-regulated at P2 in both lines (Fig. 6A, Dataset S5). This gene is closely related to the Arabidopsis gene encoding SHORTROOT (SHR) (Huang et al., 2015), which together with another GRAS-domain TF, SCARECROW (SCR), regulates the duration of cell proliferation in leaves (Dhondt et al., 2010). Moreover, consistent with previous reports, IL2-5 and IL4-3 DEGs are enriched for E2F binding site motifs (Dataset S7, Ranjan et al., 2016). E2F transcription factors act downstream of SHR and SCR to regulate progression through the S-phase of the cell cycle (Dhondt et al., 2010). These data support the notion that the extent and/or duration of cell proliferation underpin increased thickness in these lines. Another set of DEGs that distinguish the thick ILs and the Sp parent from domesticated tomato include three genes with predicted functions in regulating the cell cycle and cell expansion activities: a LIM-domain protein (Solyc04g077780), a JAZ1 TF (Solyc12g009220), and a GA 20-oxidase (Solyc03g006880) (Fig. 6). LIM-domain proteins have been implicated in a variety of functions including regulation of the cell cycle and organ size in Arabidopsis (Li et al., 2008). GA 20-oxidase encodes a key GA biosynthetic enzyme, which acts to promote cell elongation (Hisamatsu et al., 2005; De Lucas et al., 2008) and thus, determinacy during leaf morphogenesis of compound leaves, such as those of tomato (Hay et al., 2002). Moreover, JAZ proteins act as transcriptional repressors and are a central hub in the signaling circuit that integrates environmental cues, such as light quality, to balance growth and defense (reviewed in Hou et al., 2013). Finally, it is noteworthy to highlight that abiotic cues such as light quality and ABA have been shown to interact and modulate the activity of GA 20-ox and JAZ, and the Arabidopsis LIM-domain protein DA1, respectively, thereby establishing a conceptual means of environmental regulation of leaf thickness patterning. Taken together with higher endopolyploidy levels, the shared expression patterns for these genes between both thick ILs and the Sp parent suggests that leaf thickness results from an alteration in the trajectory of cellular events during leaf ontogeny, specifically, the duration of cell proliferation, and the timing and extent of cell expansion.

### Gene expression networks point to distinct leaf ontogeny in ILs 2-5 and 4-3

Since we observed a set of shared DEGs in lines 2-5 and 4-3, we hypothesized that general patterns of leaf ontogeny may also be shared between these lines, suggesting a core shared trajectory of leaf thickness patterning. However, we found that Dynamic Bayesian Networks of gene co-expression in ILs 2-5 and 4-3 are largely distinct (Datasets S8, S9; Fig. 7A, D).

For example, central to the organ-specific network of IL2-5 is an SBP-box domain gene, SBP 04g (SQUAMOSA promoter binding protein, Solyc04g064470) which is highly expressed throughout leaf development in IL2-5 (Fig. S6, Fig. 7A,B). SBP transcription factors regulate various aspects of plant growth by controlling the rate and timing of developmental events, including leaf initiation rate (reviewed in Preston and Hileman, 2013). Further, the promoters of IL2-5 DEGs are enriched for SBP motifs (Dataset S7) supporting the central role of this group of transcription factors during IL2-5 leaf ontogeny. Interestingly, GO terms for “photosynthesis” and “translation” are enriched among P4 up-regulated genes. This observation suggests that processes associated with cell differentiation (i.e. photosynthetic gene function and protein translation) are precociously activated in IL2-5 relative to domesticated tomato and supports a hypothesis whereby the overall schedule of leaf developmental events may be hastened in IL2-5.

In contrast, a central node in the IL4-3 co-expression network is a BEL1-like 04g (Solyc04g080780). BEL1-like homeodomain proteins interact with class I KNOX transcription factors to pattern the SAM and lateral organs, including leaf complexity (Kimura et al., 2008; Hay and Tsiantis, 2010) and the extent of lobing and serrations (Kumar et al., 2007). Like *S. pennellii*, IL4-3 leaflets have significantly smoother margins (fewer serrations) than M82, as reflected in increased circularity (Fig. 4B; Holtan and Hake, 2003; Chitwood et al., 2013).

These distinct dynamic patterns of leaf ontogeny that each IL shares with the desert-adapted parent may reflect aspects of leaf development unrelated to the patterning of leaf thickness, such as the patterning of leaf complexity and leaflet shape in IL4-3. Alternatively, it is also possible that the core mechanism of leaf thickness patterning is achieved by regulation of the timing and extent of cellular activities, such as the balance between cell proliferation and the onset of cell expansion and endoreduplication, with a number of potential molecular networks needed to accomplish these roles. An observation supporting this model is the fact that IL2-5 and IL4-3 have non-overlapping sets of cell cycle related DEGs. This hypothesis is consistent with the additive phenotypes of IL2-5/IL4-3 double homozygotes (Fig. 4, Fig. S4), whereby IL-specific regulators may converge on a common set of targets to regulate cell size and shape, and ultimately leaf thickness.

## Materials and Methods

### Plant material and growth conditions

Seeds for 76 *S. pennellii* introgression lines (LA4028-LA4103; Eshed and Zamir, 1995) and the *S. lycopersicum* domesticated variety M82 (LA3475) were obtained either from Dr. Neelima Sinha (University of California, Davis) or from the Tomato Genetics Resource Center (University of California, Davis). All seeds were treated with 50% bleach for 3 min, rinsed with water and germinated in Phytatrays (P1552, Sigma-Aldrich). Seeds were left in the dark for 3 days, followed by 3 days in light, and finally transferred to greenhouse conditions in 50-plug trays. Hardened plants were transplanted to field conditions at the Bradford Research Station in Columbia, MO (May 21, 2014) with 3 m between rows and about 1 m spacing between plants within rows. A non-experimental M82 plant was placed at both ends of each row, and an entire row was placed at each end of the field to reduce border effects on experimental plants. The final design had 15 blocks, each consisting of 4 rows with 20 plants per row. Each of the 76 ILs and 2 experimental M82 plants were randomized within each block. IL6-2 was excluded from final analyses due to seed stock contamination. For the analysis of leaf primordia by confocal microscopy and RNA-Seq, IL2-5, IL4-3, M82, and *S. pennellii* seeds were germinated as above and transferred to pots in controlled growth chamber conditions: irradiance at 400 μmol/m^2^/s, 23 °C, 14-hour days. Growth conditions for the drought phenotyping experiment were irradiance of 200 μmol/m^2^/s at a daytime temperature of 22 °C and 18 °C at night.

### Whole-plant phenotyping under drought

The LemnaTec Scanalyzer plant phenotyping facility at the Donald Danforth Plant Science Center (LemnaTec GmbH, Aachen, Germany) was used to phenotype approximately 3-week old *S. lycopersicum* and *S. pennellii* plants (n = 8/genotype) subjected to one of three watering regimes: 40 % field capacity, 20 % field capacity, and no watering (0 % field capacity). Top view images of each plant taken every second night over 16 nights were analyzed using custom pipelines in Lemna Launcher (LemnaTec software) to extract total plant pixel area (a proxy for biomass).

### Trait measurements

After flowering (July 2014), four fully expanded adult leaves were harvested from each plant; the adaxial (upper) surfaces of distal lateral leaflets harvested from the left side of the rachis were scanned with a flatbed scanner to obtain raw JPG files. The middle portion of each leaflet was then attached on a custom-build dual confocal profilometer device (Fig. S1) and the thickness of each leaflet was measured across the leaflet surface at a resolution of 1 mm^2^. Median thickness was calculated across each leaflet using values in the range (0 mm < thickness < 2 mm) and these median values were averaged across four leaflets per plant to arrive at a single robust metric of leaf thickness. Finally, entire leaflets were dried and their dry mass used to calculate leaf mass per area (LMA) for each leaflet. Leaflet outline scans were processed using custom macros in Image J (Abramoff et al., 2004) to segment individual leaflets and to threshold and binarize each leaflet image. Shape descriptors area, aspect ratio, roundness, circularity, and solidity (described in detail in Chitwood et al., 2013) were extracted from binary images. Additionally, elliptical Fourier descriptors (EFDs) for leaflet outlines were determined using SHAPE (Iwata and Ukai, 2002). For this analysis 20 harmonics with 4 coefficients each were used to derive principal components (PC) that describe major trends in the shape data.

### Elemental profiling (ionomics)

Distal lateral leaflets of fully expanded young (Y) and old (O) leaves of the same plants as above were collected from five individuals of each genotype. Whole leaflets were weighed and digested in nitric acid at 100 °C for 3 hours. Elemental concentrations were measured using an inductively coupled plasma mass spectrometer (ICP-MS, Elan DRC-e, Perkin Elmer) following the procedure described in Ziegler et al. (2012). Instrument reported concentrations were corrected for losses during sample preparation and changes in instrument response during analysis using Yttrium and Indium internal standards and a matrix-matched control run every tenth sample. Final concentrations were normalized to sample weight and reported in mg analyte per kilogram tissue.

### Statistical analyses and data visualization

All statistical analysis and visualization was carried out using R packages (R Core Team, 2013). QTL were identified using the mixed effect linear model packages lme4 (Bates et al., 2014) and lmerTest (Kuznetsova et al., 2015) with M82 as intercept, IL genotype as a fixed effect, and field position attributes (block, row, and column) as random effects. Only effects with significant variance (p < 0.05) were included in the final models. For elemental composition data, leaf age (“young” and “old”) was also included as a random effect unless the variance due to age was the greatest source of variance; in such cases, young and old samples were analyzed separately. Heritability values represent the relative proportion of variance due to genotype. For the quantification of organ volume parameters and photosynthesis measurements, linear models were used to test the effect of genotype. All plots were generated with the package ggplot2 (Wickham, 2009).

### Trait correlations and hierarchical clustering

For trait correlation analyses we included all traits reported in this study and measured on the same set of field-grown IL individuals (leaf thickness, LMA, leaflet shape traits, elemental profiles). We also included several sets of meta-data detailed in Dataset S3, including DEV (developmental), MOR (morphological), MET (fruit pericarp metabolite content), ENZ (enzyme activity), and SED (seed metabolite content) related traits (from Chitwood et al., 2013 and references within). Spearman correlation coefficients (rho) were calculated between each pair of traits using the rcorr function in Hmisc (Harrell et al., 2015) and p-values for the correlations were corrected for False Discovery Rate using Benjamini Hochberg (Dataset S4). Hierarchical clustering and visualization of significant correlation (q < 0.05) of leaf thickness and LMA were clustered (hierarchical “ward” algorithm) and visualized using pheatmap (Kolde, 2015).

### Estimation of nuclear size profiles by flow cytometry

Distal lateral leaflets were harvested from the 7^th^ leaf of greenhouse-grown 6-week-old plants and immediately chopped in 1 mL of ice-cold buffer LB01 as in Doležel et al. (2008). The resulting fine homogenate was filtered through a 30 um Partec CellTrics filter (5004-004-2326) and incubated with 50 ug/mL propidium iodide (Thermo Fisher, P21493) and 50 ug/mL RNase A (Qiagen, 19101) for 20 min on ice. Fluorescence scatter data was collected without gating using a BD Acuri CS6 instrument (BD Biosciences). Plots of event count as a function of fluorescence area were used to estimate the proportion of nuclei of sizes corresponding to 2C, 4C, and 8C in each genotype.

### Confocal microscopy, 3D-reconstructions, and organ volume quantification

For mature leaf cross-sections, field-grown leaves were fixed in FAA (4 % formaldehyde, 5 % glacial acetic acid, 50 % ethanol), vacuum infiltrated, dehydrated through an ethanol series, rehydrated to 100 % water, stained in 0.002 % propidium iodide (Thermo Fisher, P21493) for 2 hours, dehydrated to 100 % ethanol, and finally cleared in 100 % methyl salicylate (Sigma, M6752) for 7 days. Hand-sections were visualized with a Leica SP8 laser scanning confocal microscope using white light laser excitation at 514 nm with a 20X objective. Two partially overlapping images were captured for each cross-section and merged into a single image using the “Photomerge” function in Adobe Photoshop CC 2014 (Adobe Systems Incorporated). For the quantification of P3 leaf primordium dimensions, shoot apices (shoot apical meristem and P1-P4) of 14 day-old seedlings grown in controlled conditions were excised, fixed, processed, and stained as detailed for leaf cross sections above. Confocal stacks were obtained at software-optimized intervals, and exported as TIFF files. Raw stack files were imported into MorphoGraphX (Reulle et al., 2015). After Gaussian filtering, the marching cubes surface reconstruction function was used (cube size = 5 μm and threshold = 7,000). The resulting surface mesh was smoothed and subdivided twice and exported as a PLY file. To minimize the effects of trichomes on P3 volume, all meshes were trimmed in MeshLab (Cignoni et al., 2008). Volume, length, and diameter of processed P3 meshes were calculated using custom scripts in MatLab (MathWorks, Inc.). Briefly, first, we detected the boundary of each hole and calculated its centroid point. We connected boundary points of each hole to its centroid and filled the triangle faces. After filling all the holes, 3D mesh represents the closed surface. Then we calculated the volume based on the divergence theorem, which makes use of the fact that the inside fluid expansion equals the flux 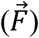 of the fluid out of the surface(S).When the flux is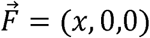 the volume is 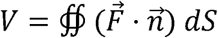, where 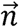 is normal vector. Thus, for each triangle, we computed the normal vector 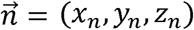 (*x*_*n*_, *y*_*n*_, *z*_*n*_), the area A, and the centroid point *P*=(*x*_*p*_, *y*_*p*_, *z*_*p*_),. The volume V is the summation of *Ax*_*n*_*x*_*p*_ for all triangles. To estimate organ arch length we made use of the fact that the Laplace-Beltrami eigenfunctions are deformation invariant shape descriptor (Rustamov, 2007). We thus employed its first eigenfunction, which is associated with the smallest positive eigenvalue and discretized the eigenfunction values into 50 sets to compute the centroid point to each set. We fit a cubic function by fixing two end-point constraints to those centroid points to get a smooth principle median axis. Note that the two end points were manually adjusted to correct for artifacts. The length of this axis is used to quantify the length of the organ. Finally, we calculated mean organ diameter as 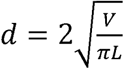.

### RNA-Seq library preparation and sequencing

Apices of fourteen day-old IL2-5, IL4-3, M82, and *S. pennellii* (Sp) plants grown in a randomized design under controlled growth conditions were hand-dissected under a dissecting microscope to separate plastochrons P4, P3, P2, and P1+SAM organs corresponding approximately to leaves L5 – L8. For Sp plants we were not able to separate P2 primordia from the apex and so we obtained P4, P3, and SAM+P1+P2 samples. Dissected organs were removed from the apex in less than 60 seconds and immediately fixed in 100 % ice-cold acetone to preserve the integrity of RNA in the sample. Each biological replicate is a pool of 10 individuals, and a total of 5 biological the surface (). When the flux is replicates were obtained for each genotype/organ combination. RNA was extracted using PicoPure RNA Isolation Kit (Thermo Fisher Scientific, MA, USA) according to the manufacturer’s protocol with the optional on-column DNase treatment. RNA integrity (RIN) was assessed by running all samples on an Agilent RNA 6000 Pico chip (Agilent Technologies, CA, USA) and three biological replicates with RIN > 7.0 were selected for further processing. Double stranded cDNA amplified using Clontech SMARTer PCR cDNA synthesis kit (634926, TaKaRa Bio USA) was fragmented for 15 min using Fragmentase (M0348, New England Biolabs) and processed into Illumina sequencing libraries as follows: the ends of 1.5X AMPure XP bead (A63880, Agencourt) purified fragmented DNA was repaired with End Repair Enzyme Mix (E6050 New England Biolabs) and Klenow DNA Polymerase (M0210, NEB), followed by dA-tailing using Klenow 3’-5’ exonuclease (M0212, NEB). The Illumina TruSeq universal adapter dimer was ligated to library fragments with rapid T4 DNA Ligase (L6030-HC-L, Enzymatics) followed by 3 rounds of 1X AMPure XP bead purification to remove unligated adapter. Finally, libraries were enriched and indexed by PCR using Phusion HiFi Polymerase mix (M0531, NEB). Illumina libraries were quantified using a nanodrop, pooled to a final concentration of 20 nM and sequenced as single end 100 bp reads on Illumina HiSeq2500 at the Washington University in St. Louis School of Medicine Genome Technology Access Center (https://gtac.wustl.edu/).

### RNA-Seq data analysis

Adapters and low quality bases were removed using Trimmomatic (Bolger et al., 2014) with default parameters. Trimmed reads were mapped to the ITAG2.3 *Solanum lycopersicum* genome (https://solgenomics.net/organism/Solanum_lycopersicum/genome; The Tomato Genome Consortium, 2012) using bowtie2 (Langmead and Salzberg, 2012) to obtain SAM files. After sorting and indexing of SAM files, BAM files files were generated using samtools commands (Li and Handsaker et al., 2009). The BEDtools multicov tool (Quinlan and Hall, 2010) was then used to obtain read counts per annotated gene for each sample. Subsequent analysis was done with the R package edgeR (Robinson et al., 2010). After normalization for library size 20,231 genes with at least one count per million reads across three samples were retained for further analysis. Lists of Differentially Expressed Genes (DEGs) were generated between pairwise sample combinations with q-value <0.05. For IL2-5 and IL4-3 at P1, P3, and P4 stages, we identified genes that are differentially expressed relative to M82 in both the IL and the Sp parent to interrogate Sp-like changes in gene expression in the ILs. For P2, the list of DEG in each IL reflects changes relative to M82 only (Dataset S5).

### Gene Ontology, Mapman, and promoter motif enrichment analyses

Lists of IL organ-specific DEGs were interrogated for enrichment of Gene Ontology terms using agriGO (http://bioinfo.cau.edu.cn/agriGO/; Du et al., 2010) with default parameters (Fisher’s exact significance test and Yekutieli FDR adjustment at q < 0.05). We further divided DEG gene lists into IL up-regulated and down-regulated genes and report significant terms in Dataset S6. We tested IL organ-specific DEGs for enrichment of annotated promoter motifs using a custom R script (Dr. Julin Maloof). Briefly, functions in the Bioconductor Biostrings package (Pages et al., 2016) were implemented to count the frequency of 100 known motifs in the promoters of DEGs (1000 bp upstream sequence) and calculate p-values for enrichment based on these counts. We report exact matches of known motifs and motifs with up to 1 mismatch in IL up-regulated and down-regulated organ-specific gene sets (Dataset S7).

### IL organ-specific gene network inference

To infer IL organ-specific networks (Fig. 7A-D, Dataset S8), we selected DEGs between IL2-5/M82 (IL4-3/M82) and Sp/M82 for each organ (P1, P3, P4) (q value < 0.05). Since co-expression analysis can inform the likelihood that genes interact, or participate in the same functional pathway, the selected genes for each IL (IL2-5 or IL4-3) and each organ were clustered based on their co-expression across genotypes. To perform clustering, the Silhouette index (Rousseeuw, 1987) followed by K-means (MacQueen, 1967) were applied. After clustering, networks were inferred as in de Luis Balaguer et al. (2017). Briefly, for each DEG, we identified a set of potential regulators and measured the likelihood of gene-target regulation using a Bayesian Dirichlet equivalence uniform (Boutine, 1991). Genes that had the highest value of the Bayesian Dirichlet equivalence uniform were chosen as regulators, and of these only transcription factors (as annotated by Suresh et al., 2014) were further considered as regulatory (source) nodes. To obtain the final IL2-5 and IL4-3 organ-specific networks, the networks for each cluster were connected. For this, we found regulations among the cluster hubs (node of each individual network with the largest degree of edges leaving the node) by using the same Bayesian Dirichlet equivalence uniform metric. In addition, we implemented a score to estimate whether the inferred interactions were activations or repressions. The score was calculated for each edge and it measured the ratio between i) the conditional probability that a gene is expressed given that its regulator was expressed in the prior time point, and ii) the conditional probability that a gene is expressed given that its regulator was not expressed in the prior time point. If the first conditional probability is larger than the second one, then the parent was found to be an activator and *vice versa*. In the case of a tie, the edge was found to have an undetermined sign. Networks for each organ were jointly visualized in Cytoscape (Shannon et al., 2003).

### Dynamic IL network construction

To infer dynamic IL networks (Fig. 7E-F, Dataset S9), we selected DEGs between IL2-5/M82 or IL4-3/M82 and Sp/M82 for each organ (P1, P3, P4) (q value < 0.05 or (FC > 2.0 and q value < 0.2)). All DEG in the IL2-5 or IL4-3 were clustered in four groups, corresponding to the four developmental stages: each gene was assigned to the developmental stage where it showed the maximum expression. A network was then inferred for each developmental stage as described for the IL organ-specific networks. To ensure that all potential regulators of each gene were considered, genes from the preceding developmental stage were included in the inference of the network of each developmental stage. The final network for each IL was visualized in Cytoscape (Shannon et al., 2003).

## Accession Numbers

An NCBI SRA accession number will be provided upon publication.

## Supporting Data

**Supplemental Figure S1.** Dual confocal profilometer device used to measure leaf thickness.

**Supplemental Figure S2.** Comparison of leaf thickness of select ILs as a function of shoot position and field vs. greenhouse conditions.

**Supplemental Figure S3.** Representative leaf cross-sections and flow cytometry of leaf 6/7 for 10 ILs harboring leaf thickness QTLs grown in greenhouse conditions.

**Supplemental Figure S4.** Mean dimensions of palisade and spongy mesophyll cell layers in select thick leaf ILs. Representative flow cytometry histograms of leaf 7 and post-flowering leaves from each genotype.

**Supplemental Figure S5.** Representative shoot apex reconstructions highlighting the appearance of early and late stage leaf primordia for each genotype in Fig. 4

**Supplemental Figure S6.** Summary of differentially expressed genes in IL2-5 and IL4-3.

**Supplemental Figure S7.** Expression profiles of differentially expressed putative transcription factors in ILs 2-5 and 4-3.

**Supplemental Figure S8**. Summary of enriched promoter motifs among differentially expressed genes in ILs 2-5 and 4-3.

**Supplemental Dataset S1.** Trait value estimates and heritability for leaf thickness, LMA, and leaflet shape across the IL panel.

**Supplemental Dataset S2.** Trait value estimates and heritability for elemental concentration across the IL panel.

**Supplemental Dataset S3.** Summary of all measured and meta-data traits used in correlation matrix.

**Supplemental Dataset S4.** Pairwise trait correlation matrix including significance values.

**Supplemental Dataset S5.** List of differentially expressed genes (q < 0.05) in each organ (P1 – P4) for the comparison: (M82/IL) overlapping with (M82/*S. pennellii*).

**Supplemental Dataset S6.** List of significantly enriched (q < 0.05) Gene Ontology (GO) terms for gene sets listed in Dataset S5.

**Supplemental Dataset S7.** List of enriched (q < 0.05) promoter motifs for gene sets in Dataset S5.

**Supplemental Dataset S8.** List of organ-specific (P1, P3, P4) gene interactions for IL2-5 and IL4-3.

**Supplemental Dataset S9.** List of dynamic gene interactions for IL2-5 and IL4-3.

## Acknowledgments

*S. pennellii* introgression line panel seeds were provided by Dr. Neelima Sinha (University of California, Davis) and the Tomato Genetics Resource Center (University of California, Davis). We would like to thank Dr. Ivan Baxter and Dr. Greg Ziegler (Danforth Plant Science Center) for generating elemental profile data, and Dr. Julin Maloof (University of California, Davis) for sharing custom promoter enrichment analysis scripts. We acknowledge the advice and assistance of Dr. Noah Fahlgren, Dr. Malia Gehan, and Melinda Darnell (Danforth Plant Science Center) with drought phenotyping experiments. We thank Dr. Elizabeth Kellogg for insightful discussions and comments on the manuscript. This work was supported by funds from the Donald Danforth Plant Science Center. RS is supported by an NSF CAREER grant (MCB-1453130). MHF is supported by an NSF NPGI post-doctoral fellowship (IOS-1523668).

